# Modulation of gut microbiota through *in vitro* exposure to a traditional fermented food in the context of pathogen invasion

**DOI:** 10.1101/2025.03.25.645194

**Authors:** Murambiwa Nyati, Oscar van Mastrigt, Elise F. Talsma, John Shindano, Bas J. Zwaan, Sijmen Schoustra

## Abstract

The gut microbiota consisting of bacteria, fungi, archea, and viruses and plays a role as a barrier against pathogen invasion to promote intestinal health and preventing organismal infections. This study examined the potential protective properties of Mabisi, a widely consumed traditional fermented dairy product in Zambia, against pathogen invasion in the gut. Using the *in-vitro* static INFOGEST model, we challenged the gut microbiota from a pooled stool sample pf six Zambian children) and previously exposed to Mabisi by adding model pathogens. We hypothesized that pathogen invasion is suppressed in *in-vitro* gut microbiota treated with Mabisi due to its ability to modulate the gut microbiota composition. Our microbiota challenge treatments were Mabisi, fructooligosaccharides (FOS) as a positive control, sterile water as a negative control, and with and without pathogens. Out of the two strains of pathogens added to the gut microbiota, *Escherichia coli* established in FOS treated gut microbiota, however not in Mabisi and sterile water treatments. Furthermore, we show that Mabisi modulated gut microbiota by increasing *Pediococcus*, a commensal bacteria and increased production of short chain fatty acids including acetate, propionate, lactate, isobutyrate, formate, and succinate in Mabisi treated gut microbiota compared to both FOS and sterile water treated gut microbiota. This study contributes to evidence of the beneficial role of fermented foods such as Mabisi through the prevention of pathogen invasion of the gut microbiota in children.

**Importance:** This study provides evidence supporting the protective role of Mabisi, a traditional fermented dairy product, in preventing pathogen invasion in the gut microbiota of children. By using an in-vitro approach, we demonstrate that Mabisi treatment suppresses the establishment of Escherichia coli, a key pathogen, unlike fructooligosaccharides (FOS), a well-known prebiotic. Additionally, Mabisi modulates gut microbiota composition by increasing beneficial bacteria such as *Pediococcus* and enhancing short-chain fatty acid (SCFA) production, which is essential for gut health.

These findings highlight the potential of traditional fermented foods as affordable, accessible dietary interventions for improving gut health and preventing infections, particularly in low-resource settings like Zambia.

## 1.0 Introduction

The human gut microbiota is a complex ecosystem comprising bacteria, fungi, archaea, and viruses, each contributing to maintaining host health. Among these, certain bacterial groups particularly support gut health and immune function of the host. A healthy gut microbiota consists of bacterial community that is biased towards the dominance of commensal bacterial species. Commensal bacterial species live in harmony with host and may improve digestion and strengthen immune functions. In contrast to pathogenic species, commensal species do not cause disease under normal conditions. Depending on bacterial taxa composition, the gut microbiota serves as a barrier against pathogen invasion.

Environmental factors are known to shape gut microbiota, some promote healthy gut microbiota to prevent invasion of pathogenic bacteria species^1^. Environmental factors that may impact gut microbiota include nutrition, antibiotic use, exposure to environmental microbes, pollution and toxins, water quality, and hygiene and sanitation. A nutritional factor that is the focus of this study, is the consumption of traditional fermented foods (TTF) as they have the potential to alter gut microbiota to a host’s advantage^2^. Indeed, TTFs can modulate the gut microbiota modulation such that it increases the resilience to withstand invasion by pathogenic bacterial species^3^. In the sub-Saharan Africa region, consumption of dairy and cereal based TFF such as Masi, Munkoyo, Chibwantu, Mabisi, Ogi, and Bushera is common. TFF are commonly produced through natural fermentation making it a rich source of environmental microbes and metabolites. Environmental microbes act on carbohydrates to produce metabolites such as short chain fatty acids (SCFA) which are beneficial for host health^4^.

Studying pathogens invasion of gut microbiota using *in-vivo* approaches in humans or animals is challenging; therefore, *in-vitro* digestion models are useful alternatives. *In - vitro* digestion models range from simple or basic batch models based on test tubes to complex models running on sophisticated programs. For instance, the INFOGEST static digestion is a simple model offers efficient and high replication experimentation leading to reproducible outcomes ^5–8^. Furthermore, the model provides a cost effective solution to studying the effect of nutrition on gut microbiota, which is especially welcome in low-income settings.

In this study, we investigate whether the gut microbiota is more resilient to the invasion of known pathogenic bacterial species after exposure to the Zambian TFF, Mabisi. The microbiota was derived from a pool of stool samples taken from six children from rural Zambia. Specifically, we hypothesize that invasion by pathogenic bacterial species is suppressed when gut microbiota is exposed to Mabisi.. Mabisi has been reported to harbour a diverse community of lactic and acetic acid producing bacteria ^9^.

## 2.0 Methods and materials

### Experiment setup

We conducted this experiment following the *in vitro* INFOGEST static and colonic digestion protocols as suggested by Minekus, Brodkorb and Perez-Burillo^10–12^. These protocols are validated and used to study food ingredients and its modulations of the gut microbiota^8,13^.

We used three treatments, 1) Mabisi, 2) Fructooligosaccharides (FOS) as a positive control, and 3) Sterile water as a negative control. We digested treatments following a two-stage digestion procedure that simulates *in vivo* human digestion. Stage one digestion involves digestion of treatments following conditions of the oral, gastric, and small intestine phases (see supplement Figure 1). In the second stage of the experiment, colon digestion contents were prepared using digests of Mabisi, FOS, and Sterile water digests combined with a pooled stool slurry with and without pathogens (see Figure 1). The colon digestion contents were anaerobically incubated for 24hrs. Each treatment, Mabisi (exposed with or without pathogens), FOS (with or without pathogens), and sterile water (with or without pathogens) was replicated eight times.

**Figure 1.**
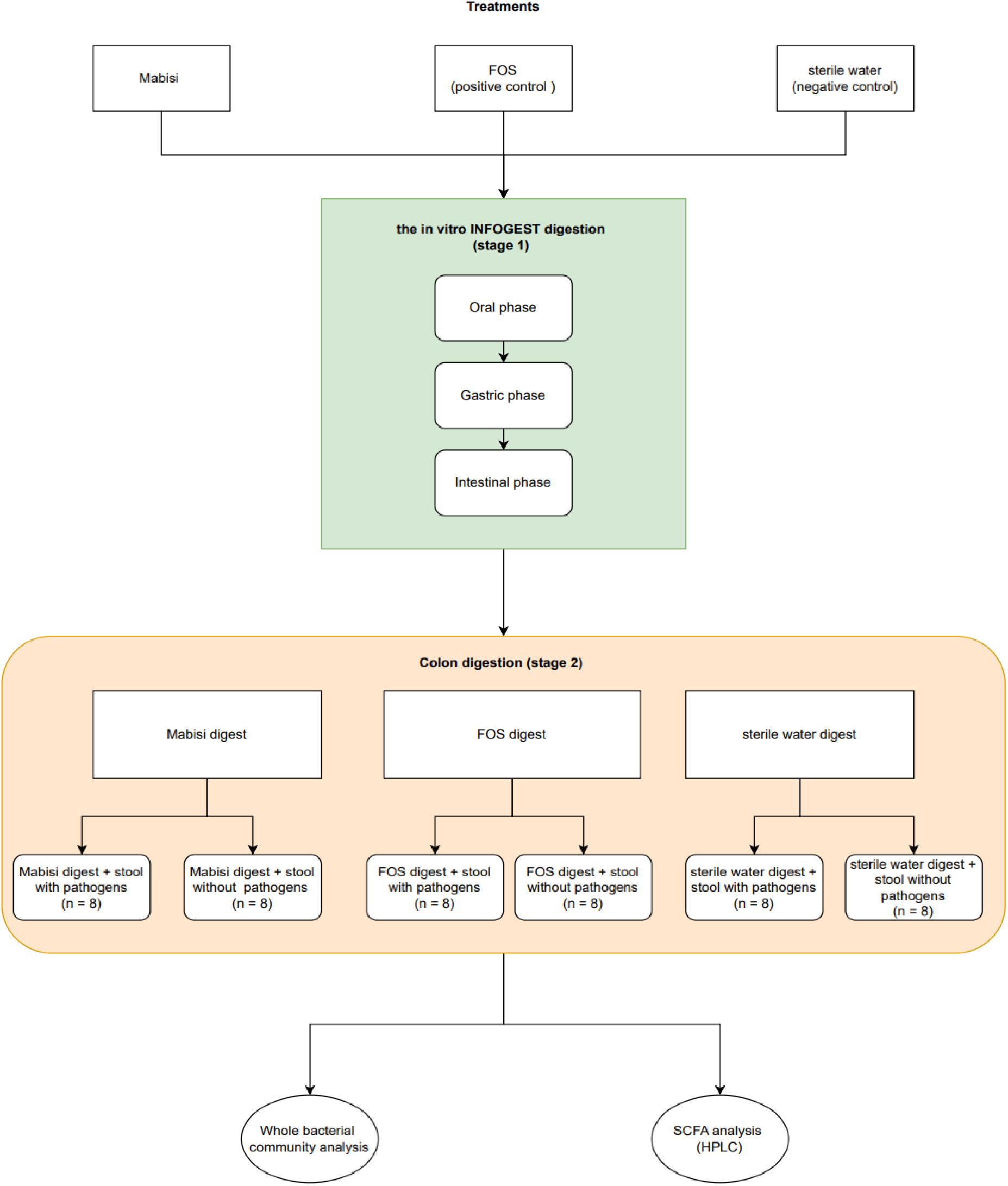
A flowchart showing *in-vitro* INFOGEST static digestion of experiment inputs (Mabisi, FOS, and sterile water). Later in the chart, Mabisi, FOS, and sterile water digests are further colon digested with pooled stool slurry with and without model pathogens. At the end of the experiment, colon digestion units are sampled for whole bacterial community and SCFA analyses.

Donors of stool were healthy children consisting of three females and three males all aged between 6 to 12 months, that were regular consumers of Mabisi and did not use antibiotics in the last 30 days. Consent was obtained from parents or caregivers prior to the study. Pathogen strains *Escherichia coli* K12 DSM498 and *Listeria innocua* ATCC 33090 DSM20649 for invasion of Mabisi, FOS, and sterile water treated colon digestion units (gut microbiota) were obtained from the department of Food Microbiology, Wageningen University and Research, the Netherlands.

### Chemicals and reagents

Soluble potato starch (S2630 Sigma-Aldrich), 3,5-dinitrosalicylic acid (DNS) (210-204-3 Sigma-Aldrich), sodium phosphate buffer (P4922 Sigma-Aldrich), sodium potassium tartrate tetrahydrate (S2377 Sigma-Aldrich), maltose standard (M5885 Sigma-Aldrich), Hydrochloric acid (H1758 Sigma-Aldrich), sodium hydroxide (S58891 Sigma-Aldrich), trichloroacetic acid (T6399 Sigma-Aldrich), sodium taurodeoxycholate solution (904236 Sigma-Aldrich), tributyrin (T8626 Sigma-Aldrich), p-toluene-sulfonyl-L-arginine methyl ester-TAME (T4626 Sigma Aldrich), bovine blood heamoglobin (H2500 Sigma-Aldrich), porcine pepsin (P6887 Sigma Aldrich), human salivary amylase (A1031, Sigma Aldrich), porcine pancreatin (P7545 Sigma Aldrich), rabbit gastric extract powder (RGE25-Lipolytech), Phosphocreatine disodium salt hydrate (ED2SC Sigma Aldrich), zinc sulfate heptahydrate (Z0251 Sigma-Aldrich), fructooligosaccharides from chicory (F8052 Sigma-Aldrich), peptone from potatoes (83059 Sigma-Aldrich) and SCFA standards (acetate, propionate, isobutyrate, butyrate, formate and lactate) all supplied by Sigma-Aldrich.

### Equipment

The following equipment were used for various procedures; Heidolph Titramex 1000 Oscillator (Hidolph, Germany), Techisch incubator(Techniekwe 18, Netherland), High Pressure Liquid Chromatography with Ultra Violet detector (Thermo Scientific, Netherlands), pH meter (Metrohm, China), Vortex machine (IKA® Genius 3, Germany).

### *In vitro* INFOGEST static digestion (stage 1)

#### Mabisi, FOS and sterile water Mabisi

We used ultra-heated full fat milk from a local supermarket in the Netherlands to prepare Mabisi. Using a previous stock of Mabisi, we inoculated 2 mL of Mabisi into ultra-heated full fat milk to produce 1000 mL of Mabisi^9^. To produce Mabisi, we incubation inoculated ultra-heated full fat milk to produce at 28°C for 48hrs. At the end of incubation we obtained a thick, white, fermented viscous liquid known as Mabisi.

#### Fructooligosaccharides

To prepare 5% FOS, we dissolved 5 g of analytical grade FOS in sterilized water to make 100 mL of 5% FOS.

#### Sterile water

Already prepared distilled and demineralized tap water was used as sterile water for this experiment.

#### Enzyme activity assay

We assessed human digestion exoenzymes for their activities to digest Mabisi and FOS. Here, we used protocols recommended by Minekus ^14^ and updated by Brodkorb ^15^.

Following protocols for salivary amylase, pepsin, lipase, trypsin and lipase in pancreatin assays, we calculated their respective activity and determined their concentrations and volumes for use. Details of concentrations and volumes of enzymes used for enzymes are mentioned later.

#### Stock chemical solutions

We prepared 1000 mL of each stock chemical solution: 0.5M KCl, 0.5M KH_2_PO_4_, 1M NaHCO_3_, 0.15M MgCl_2_(H_2_O)_6_, 0.5M (NH_4_)2CO_3_, and 0.3M CaCl_2_(H_2_O)_2_, for formulation of simulated salivary fluid (SSF), simulated gastric fluid (SGF), and simulated intestinal fluid (SIF).

#### Processing of simulated digestive fluids Simulated salivary digestion stock fluids

We prepared 1200 mL stock solution of simulated salivary fluid (SSF) using specific volumes of stock chemical solutions by mixing different chemicals of [45.3mL KCl, 11.1 mL KH_2_PO_4_, 20.4 mL NaHCO_3_, 1.5 mL MgCl_2_(H_2_O)_6_, 0.18 mL (NH_4_)_2_CO_3_] with 0.075 mL CaCl_2_(H_2_O)_2_ added immediately before use. To finalize preparation of SSF solution, we marked up to 1200ml using sterilized water.

#### Simulated gastric digestion stock fluids

For simulated gastric fluid (SGF), we mixed the following volumes of chemical solutions: [KCl (20.7 mL), KH_2_PO_4_ (2.7 mL), NaHCO_3_ (37.5 mL), MgCl_2_(H_2_O)_6_ (1.2 mL), (NH_4_)_2_CO_3_ (1.5 mL), and NaCl (35.4 mL)]. Prior to use, we added 0.015 mL of CaCl_2_(H_2_O)_2_ marked up to 1200 mL using sterile water.

#### Simulated intestinal digestion stock fluids

To prepare simulated intestinal fluid (SIF), we mixed already the following prepared stock chemical solutions: [KCl (20.4 mL), KH_2_PO_4_ (2.4 mL), NaHCO_3_ (255 mL), MgCl_2_(H_2_O)_6_ (0.9 mL), and NaCl (115.2 mL). Before use, we added 1.8 mL of CaCl_2_(H_2_O)_2_ and marked up to 1200 mL using sterile water.

### *In vitro* static INFOGEST digestion of Mabisi, FOS, and sterile water

As a first stage of the experiment, Mabisi, FOS and sterile water are separately digested following protocol 2.0. The INFOGEST digestion consists of incubation of Mabisi, FOS and sterile water using simulated digestion fluids accompanied by addition of respective phasic enzymes at oral, gastric and intestinal phases (see supplement Figure 1). Below, we describe actual procedures for each digestion phase.

#### Oral phase digestion

To prepare oral phase contents, we mixed Mabisi (5 mL) with SSF (4.5 mL) using a 50 mL centrifuge tube. We then added human salivary amylase to attain an activity of 75 U/mL. To adjust the contents pH to 7.0, we added a pre-determined volume of 0.1M NaOH. We adjusted total oral phasic content to 10 mL using sterile water and finally incubated at 37°C for 2 minutes.

#### Gastric phase digestion

To simulate normal gastric digestion, we added 9 ml SGF to 10ml oral contents obtained in the oral phase. To obtain 2000 U/ml activity of Pepsin, we added 0.7ml Pepsin dissolved in HCl. In order to obtain 60 U/ml gastric lipase activity, we added 0.3 ml of rabbit gastric extract to gastric contents. We then adjusted the pH of the solution to 3.0 using 0.1M HCl. Finally, we adjusted gastric contents by marking up to 20mls. To achieve incubation, we kept gastric contents at 37°C for 2hrs.

#### Intestinal phase digestion

To imitate small intestine digestion, we added 15ml SIF to 20ml gastric phase solution. We then added Pancreatin of 13.37mg/ml, bile salts (10mM) and adjusted pH to 6.8.

Finally, we marked up contents to 40ml prior to incubating at 37° for 2hrs.

### *In vitro* static colonic digestion of Mabisi, FOS and sterile water (stage 2)

#### Processing of small intestine digests (Mabisi, FOS and sterile water) for colon digestion

We centrifuged small intestine digests of Mabisi, FOS and sterile water at 16,000 rpm to obtain solid and supernatant fractions. For each sample, we collected supernatant volume equivalent to 10% of the solids or semi-solids for use during the colon digestion phase of the experiment. The supernatant liquid added to solids simulated the physiological fluid that transits from the small intestines to the large intestine. Specifically for this experiment, we used 0.5g of solid, and 0.62 ml of supernatant add as Mabisi, FOS and sterile water digests.

#### Processing of stock fermentation medium for colon digestion

To prepare 1000 mL of stock fermentation medium for addition in the colon digestion, we separately made 1000 mL peptone (15g/L) and 50mL reductive solutions. Reductive solution is formulated by dissolving 312mg of Cysteine and 312mg Sodium sulfide in 100 mL of distilled water. In 950 mL of peptone solution, we added 50 mL of reductive solution and autoclaved to temperature of 120°C for 2hrs. After cooling, we added 1.25 mL of filtered resazurin (0.1% w/v) and marked up to 1000ml.

#### Processing of stool slurry

To prepare stool slurry, we pooled and homogenized equal parts of the stool samples in 0.1M phosphate buffer. To make 32% (w/v) of stool slurry, we dissolved 32g of stool slurry in 100ml (100mM) phosphate buffer and adjusted of pH to 6.8. To remove bigger particles in stool slurry, we centrifuged at 550g for 5 minutes^16^. We separated the resulting stool slurry into two equal aliquots, with one part for inoculation with pathogens.

#### Description of pathogens used to inoculate stool slurry

For this experiment we used two bacteria strains *(Escherichia coli* K12 DSM498 and *Listeria innocua* ATCC 33090 DSM20649) as pathogens and potential invaders of the gut microbiota. *Escherichia coli* is a gram-negative, rod shaped facultative bacteria, and potentially pathogenic bacteria. *Escherichia coli* invades hosts by attaching itself to the mucosa tissue and effacing lesion via type III secretion system to produce specific toxins^17^. *Listeria innocua* is not a pathogen itself, but as a pathogen in lieu for *Listeria monocytogenes. Listeria monocytogenes* is a gram-positive facultative bacteria that is disease causing and can be found living in water, soils, decaying plants, and animal tissues^18,19^. It infects by invading the host cells inducing its own uptake through internalin avoiding the phagosomes and accessing intracellularly using actin-based motility.

#### Inoculation of stool slurry with pathogens

We prepared stock cultures of the pathogen strains in brain heart infusion broth. We estimated bacterial density of the stock culture using plate counts, revealing a bacterial density of approximately 5 log cfu/ml in the stock cultures. To facilitate handling during the experiment, we prepared multiple 10 mL stool slurries, with and without pathogens. Pathogen-containing slurries were prepared by adding 3 mL of pathogen stock (1.5 mL *E. coli* and 1.5 mL *L. innocua*) to 7 mL of stool slurry, while control slurries were prepared by mixing 7 mL of stool slurry with 3 mL of sterile water.

#### Colon content mixing, purging and incubation

Finally, we mixed I.5 mL of Mabisi, FOS or Sterile water digests from the small intestine phase with 7.5 ml of colon fermentation medium, 2 ml of stool slurry with or without pathogens. Subsequently, we purged the mixed colon digestion contents with nitrogen gas for 1 minute to eliminate any residual oxygen, then sealed. Lastly, we incubated on an oscillator (600 rpm) at 37°C for a period of 24 hours.

#### Sampling of colon digestion units for DNA and SCFA

Following a 24-hour incubation period, the colon digest samples underwent immediate shock treatment by immersion in ice-cold water for a duration of 5 minutes. To curtail any ongoing microbial activity, we kept the samples at −20°C. For DNA extraction, we thawed samples and centrifuged at 16,000 rpm to yield distinct phases, supernatant and solid fraction. We used the solid fraction for extraction of DNA with supernatant fraction for SCFA. Specifically, we sampled 1.5 ml of the supernatant fraction for SCFA extraction and used 250 mg of solid fraction for DNA extraction.

### DNA extraction and gut microbiota profiling

We performed bacterial DNA extraction of Mabisi, FOS, and Sterile water samples of the colon digestion phase using the DNeasy® Power Soil Pro Kit (Qiagen), adhering to the manufacturer’s guidelines^20^. Precisely, we sampled 250 mg of solid fractions of Mabisi, FOS, and Sterile water colon digestion samples and mixed with 800 µl of CD1 solution in 2 ml PowerBead Pro tubes containing fine ceramic beads supplied within the kit. To mechanically lyse bacterial cells in samples, we vortexed for 12 minutes. We then removed unwanted sample matrix through centrifugation at 15,000g for 1 minute, resulting in the collection of a supernatant enriched with DNA. Targeting the supernatant, we carefully transferred to new MB spin columns for a series of washing steps. We eluted DNA using a 90 µL elution buffer solution. To ascertain the concentration and quality of the isolated DNA, we evaluated DNA isolates using a NanoDrop™ ND-2000 and a Qubit™ 4 fluorometer (Thermo Fisher Scientific, UK).

### 16S RNA sequencing and pre-analysis processing

For identification of whole bacterial communities present in the DNA extracts, we targeted V3-V4 hypervariable region of the 16S rRNA gene amplicon. Sequencing was achieved using the NovoSeq Illumina 6000 (strategy PE250) platform at Novogene Europe (Cambridge, UK). Polymerase chain reaction (PCR) amplified the gene amplicons on 2% gel electrophoresis, using primers 341F CCTAYGGGRBGCASCAG and 806R GGACTACNNGGGTATCTAAT. Equal amounts of PCR products for each sample were pooled, end-repaired, A-tailed, and ligated with Illumina adaptors.

Paired-end reads were then generated and processed using the fast length adjustment of short reads (FLASH) software to generate 250 base pairs (bp) paired-end raw reads^21^. Subsequently, the resulting paired-end fastq files underwent merging, denoising, and filtering process to eliminate chimeric reads using the DADA2 R package^22^. This approach resulted in a dataset of high-quality, amplicon-sequenced variants (ASVs) by retaining sequences with retention criteria set at an abundance of 5% or higher.

The ASVs derived from this refined pipeline constitute a cleansed dataset facilitating the identification of bacterial taxa and the estimation of composition.

#### Tracing added model pathogens in colon digestion units

To trace and identify ASVs matching with model pathogen strains in Mabisi, FOS and Sterile water colon digestion samples, we built a standalone database containing the v3-v4 ASV sequences obtained from NovoSeq Illumina 6000 (strategy PE250) platform at Novogene Europe (Cambridge, UK). We then combined it with v3-v4 sequences of the added pathogen strains to allow for matching. Using FastTree 2 R package, we inferred evolutionary relationships of all ASVs sequences based on approximate maximum likelihood method^23^. Evolutionary relationships of ASVs matching those of model pathogen strains were mapped onto a phylogenetic tree to visualize the ASVs most likely corresponding to the added pathogen strains (see supplement Figure 2). We determined absolute abundances of ASVs (ASV 2, ASV 755, ASV 2623 and ASV 4631) closely corresponding to added pathogens. We log transformed absolute abundances to make comparisons across colon digestion treatments using box plots (see Figure 3).

### SCFA analysis

We filtered the supernatant fraction of Mabisi, FOS and sterile water colon digestion phase end products using a 0.2nm nylon filter to remove debris. Using Carrez solution (Carrez A-K4Fe(CN)6.3H2O and Carrez B-ZnSO4.7H2O) we precipitated protein from the filtered samples. To prepare Carrez solution, we mixed cold 0.25ml Carrez A solution with 0.5ml of filtered sample. Subsequently, we added cold 0.25ml Carrez B solution to precipitate protein matrix in the filtered sample. We centrifuged the resulting solution at 13000 rpm for 5 minutes to remove precipitate. We transferred precipitate free aliquot to clean HPLC standard vials (1.5ml) for SCFA extraction.

We targeted to quantify acetate, propionate, butyrate, isobutyrate, succinate, formate and lactate using a precipitate free aliquot collected in HPLC standard tubes. Our strategy for SCFA extraction followed the protocol suggested by Perez-Burillo et al^16^. High Pressure Liquid Chromatography(HPLC) with a UV detector was used to detect and quantify SCFA from samples. The full HPLC system consisted of P4000 gradient pump fitted with a vacuum degasser. Additionally, the HPLC used an auto sampler (AS3000) and a UV6000 detector (set at 210nm) all supplied by Thermo Scientific, Breda, The Netherlands^24^. We developed a calibration curve using internal standards for acetate, propionate, butyrate, isobutyrate, succinate, formate, and lactate that we prepared in a series of concentration from 0.025mM to 100mM and a blank included. Results obtained in millimolar (mM) concentration were proportional to the area under respective chromatogram. We used Chromeleon chromatography data system software^25^ to calculate actual concentration of each SCFA using peak ratio area generated using the HPLC.

### Statistical analysis

For all descriptive and inferential analysis, we used R (version 4.3.1)^26^.

We used stacked bar charts to show relative abundances of bacterial composition for Mabisi, FOS, sterile water colon digestion phase samples with and without pathogens, and analyzed relative abundances at both phylum and genus taxonomical levels. Bacterial composition relative abundance at phylum and genus levels are presented as means ± standard deviation.

Using a Phyloseq object, we calculated Bray Curtis similarity matrix (ANOSIM). We visualized similarities among Mabisi, FOS, sterile water treatments using Principal Coordinate Analysis (PCoA)^27^.

To establish dissimilarities between treatments, we performed pairwise permutation analysis of variances (PERMANOVA)^28^. Based on Benjamin Hochberg post hoc analysis, we adjusted for multiple testing. We used the adonis_pairwise function in vegan package (version 1.2.3) to generate pairwise differences^28^. Further, we identified enriched taxa in treatments using Linear discriminant analysis (LEfSe)^29^. We used 4.0 as cutoff score and false discovery rate of 5%. Welch’s ANOVA with Games Howell pairwise tests established between differences in SCFA of Mabisi, FOS, sterile water treatments^30^. We visualized SCFA relative abundance of Mabisi, FOS, sterile water treatments using PCA and violin plots.

## 3.0 Results

We report on whether Mabisi treatment affects the invasion of pathogens by comparing the gut microbiota composition with our positive and negative controls (FOS and sterile water, respectively). Further, we describe how the Mabisi treatment (relative to the controls) affects the production of short-chain fatty acids (SCFAs).

### Impact of in vitro treatment on pathogen invasion and gut microbiota species composition

Figure 2 shows the impact of Mabisi, FOS, and sterile water *in-vitro* treatments on gut microbiota, both with and without pathogens added. We tracked the added focal *Escherichia coli* and focal *Listeria innocua* strains, revealing invasion of *E. coli* under the FOS treatment. We did not detect the focal *E. coli* in the gut microbiota under the Mabisi nor sterile water treatment. Further, we did not detect *Listeria innocua* in any of the treatments (see supplement Figure 3). Exposure to Mabisi and FOS led to an increase in the relative abundance of *Lactobacillus*, *Pediococcus* and *Bifidobacterium*. Conversely, exposure to FOS led to higher relative abundance of *Enterobacter*, *Escherichia Shigella*, and other *Proteobacteria* compared to Mabisi and Sterile water treatments (also see Figure 2). ASV2, here represented as proteobacteria, increased in only pathogen added colon digestion units for FOS, however remained suppressed for Mabisi and sterile water colon digestion units exposed to pathogens (see supplement Figure 3).

**Figure 2.**
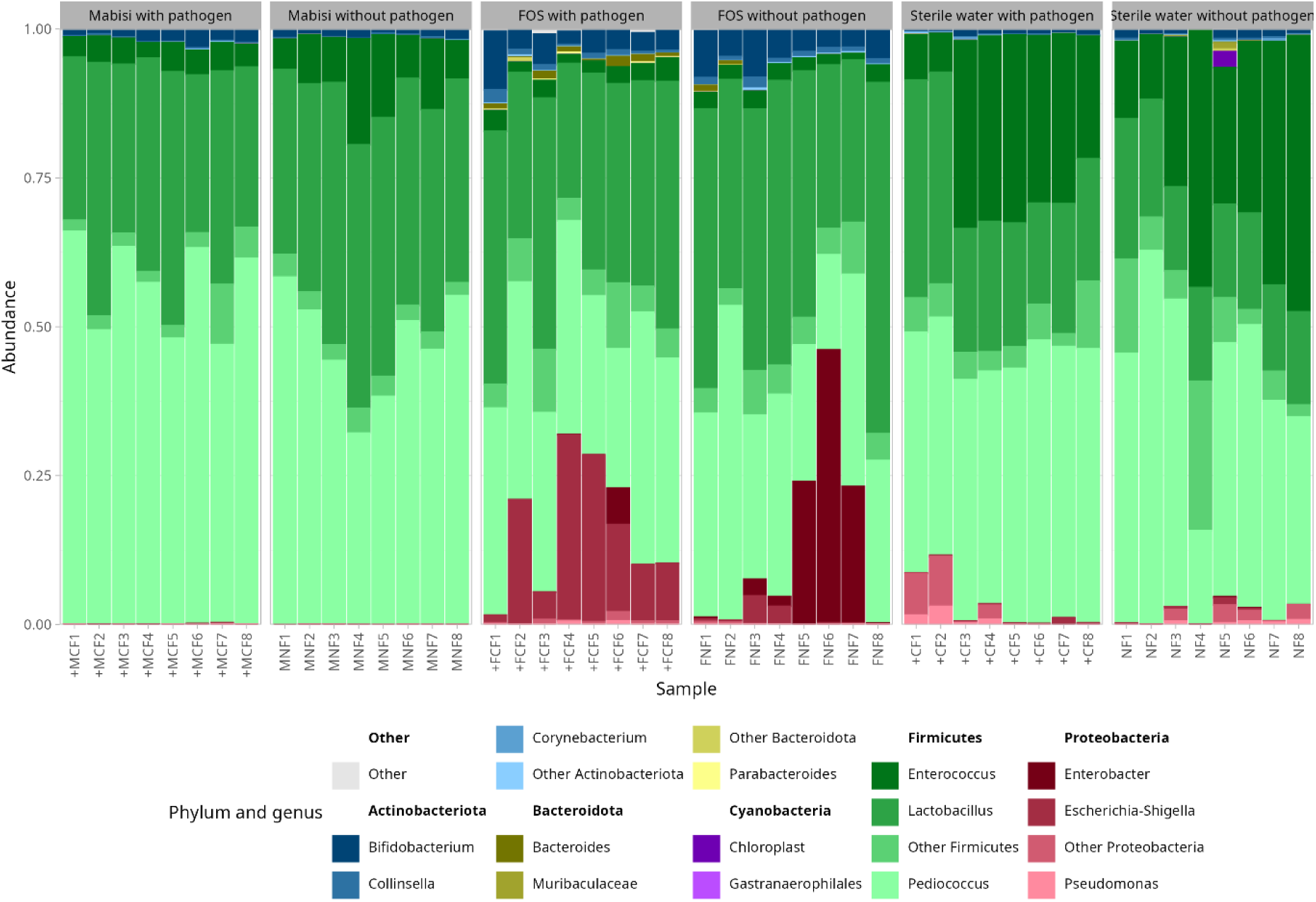
Stacked bar chart showing gut microbiota composition at phylum and genus levels of colon digestion units treated with Mabisi, FOS, and sterile water. Model pathogens, identified as *Escherichia shigella* (ASV2), are highlighted. Refer to the figure legend to navigate relative abundances by matching colour codes.

**Figure 3.**
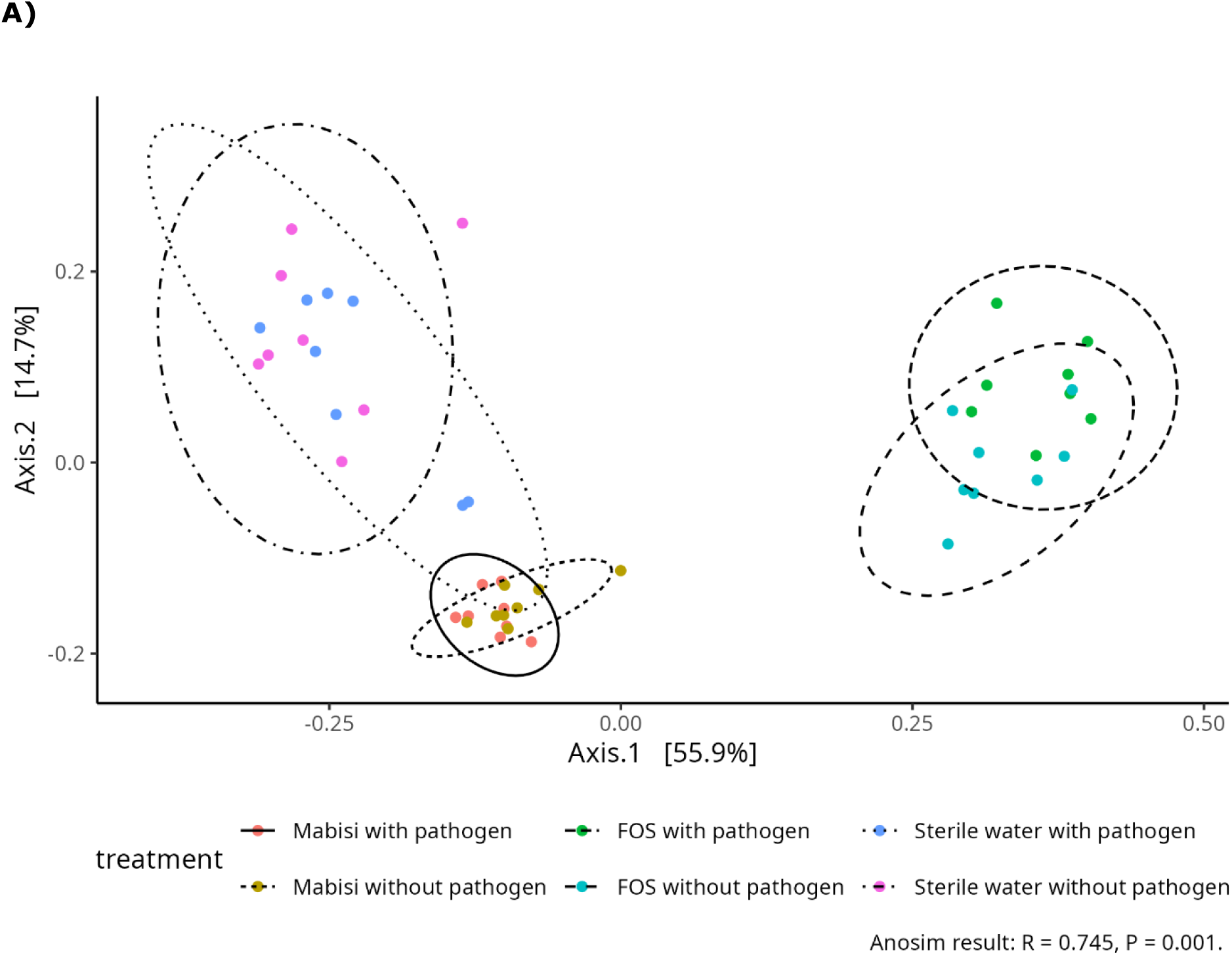
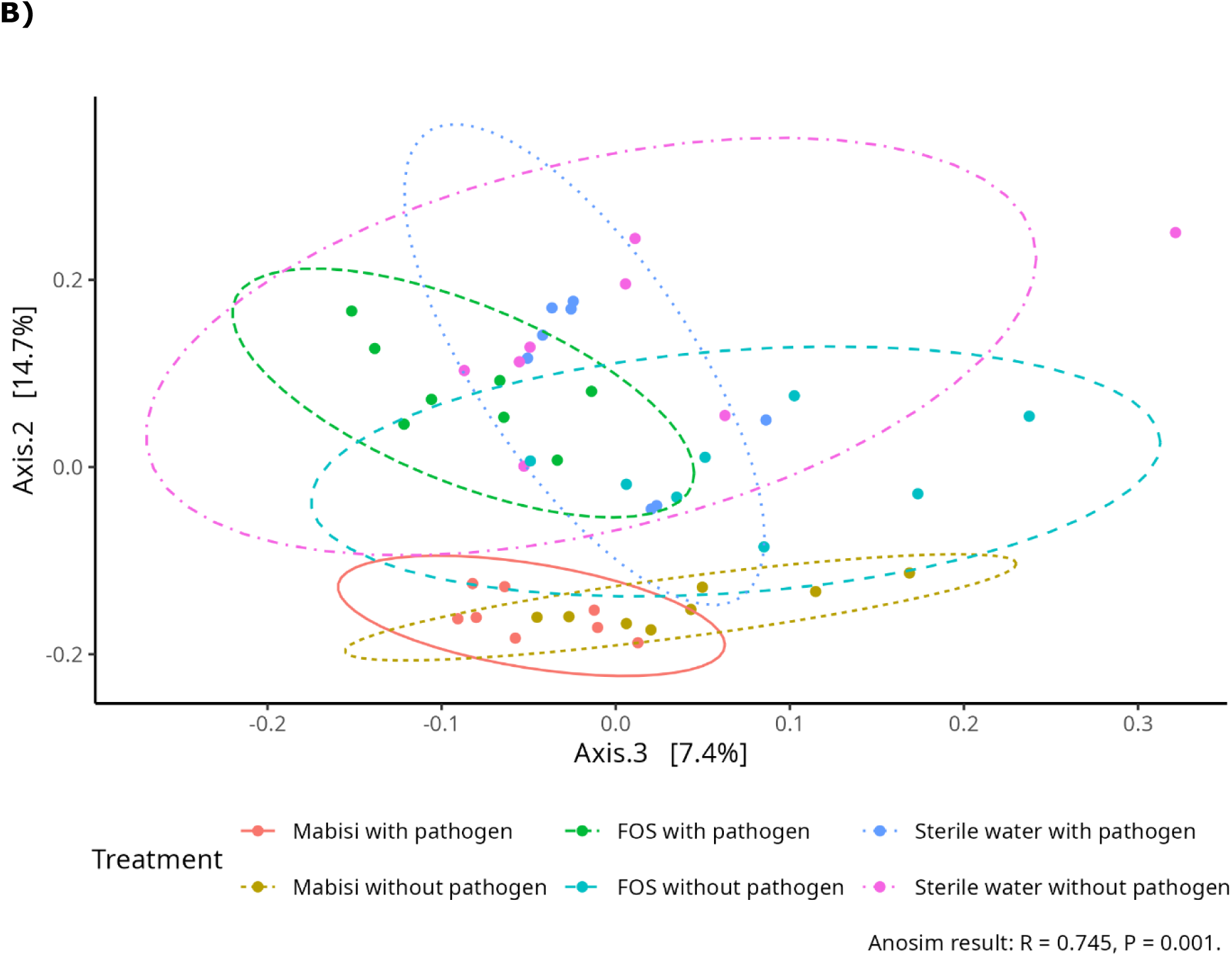
Principal coordinate analysis (PCA) of colon digestion units (with and without pathogens) treated with Mabisi, FOS and Sterile water**. A)** Axis 1 versus axis 2. Shows how the first axis separates the FOS treatments from the other treatments and the second axis the Mabisi and sterile water treatments. **B)** Axis 3 seems to separate the effect of addition of pathogens to treatments correlating to of 7.4% explained variation.

Furthermore, *Enterococcus* relative abundance in sterile water treatments is statistically significantly higher compared to both Mabisi and FOS treatments (supplementary Table 1). Principal Coordinate Analysis (PCoA) visualizes overall differences in gut microbiota composition for Mabisi, FOS, and Sterile water treatments (Figure 3A), and revealed distinct differences in gut microbiota based on treatment (Analysis of Similarities (ANOSIM); R = 0.85, P=0.001). The first axis separates the FOS treatment, the second axis the Mabisi from the sterile water treatments. We observe further cluster separation along axis 3 (Figure 3B) with pathogen addition explaining 7.4% of the total variation (also see supplement Figure 5). Specifically, pairwise comparisons revealed that the FOS treatment exposed to pathogens resulted in a different gut microbiota composition compared to not exposing to pathogen (Table 1). Pairwise comparisons of all colon digestion units are in supplement Table 2.

**Table 1:** Pairwise comparison of gut microbiota (colon digestion units) treated with Mabisi, FOS, Sterile water.

| Pairs of treatments | Sum of squares | F.Model | R2 | p.value | p adjusted |
| --- | --- | --- | --- | --- | --- |
| Mabisi with pathogen vs Mabisi without pathogen | 0.444 | 1.101 | 0.073 | 0.187 | 1.000 |
| Sterile water with pathogen vs Sterile water without pathogen | 0.511 | 1.136 | 0.075 | 0.055 | 0.831 |
| FOS with pathogen vs FOS without pathogen | 0.668 | 1.657 | 0.106 | 0.003 | 0.048 |

Using Linear Discriminant Analysis (LDA) with a threshold score of 4.0 and FDR-adjusted p-values, we identified bacterial genera that discriminate among treatments (Mabisi, FOS, and sterile water). Treatment with Mabisi increased *Pediococcus*. FOS treatment increased *Lactobacillus, Bifidobacterium, Escherichia/Shigella* (i.e. the added pathogen), and *Enterobacter*. Sterile water treatment increased *Enterococcus* and *Lactococcus* (Figure 4).

**Figure 4.**
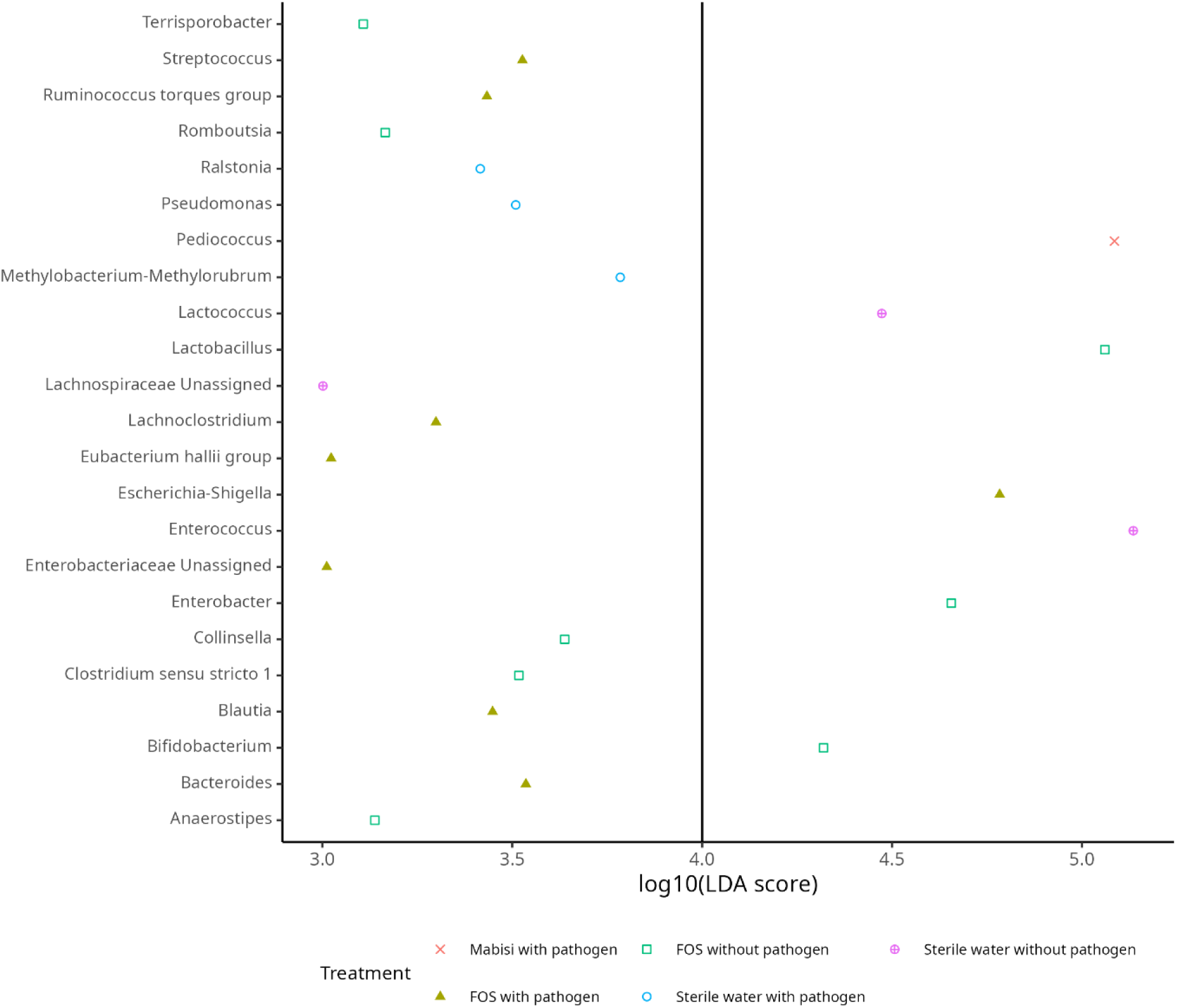
Linear discriminant analysis effect size analysis of colon digestion units after treatment with Mabisi, FOS and Sterile water.

### SCFA production in fermentation units after treatment

All Mabisi treated colon digestion units have higher concentration of acetate, isobutyrate, propionate, formate, lactate, and succinate compared to FOS and sterile water treated colon digestion units (Figure 5 & supplement Figure 7). These differences are evident with both a Welch’s ANOVA and a Game Howell pairwise test (see supplement Figure 6).

**Figure 5.**
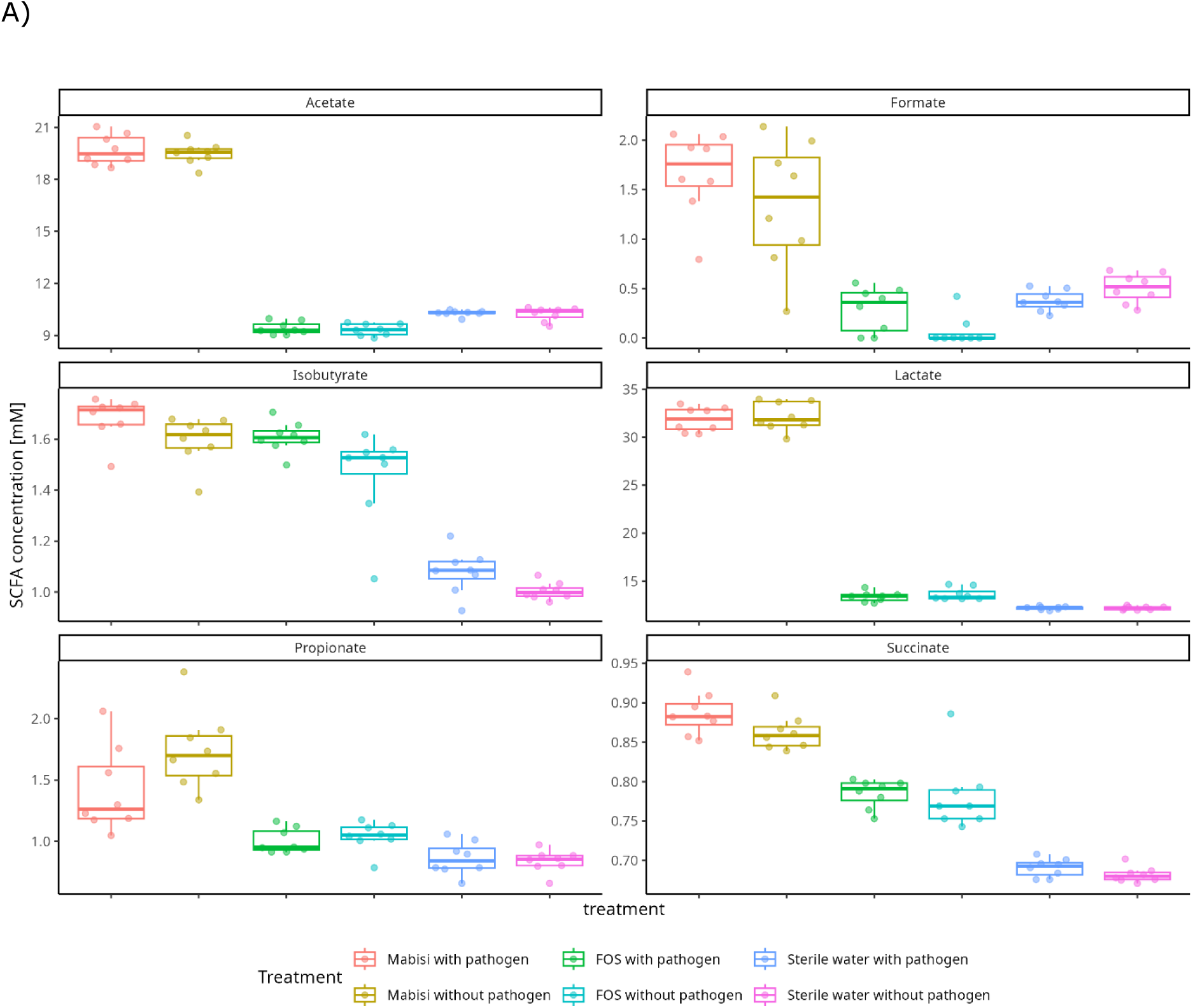
Boxplots showing concentration of SCFA in colon digestion units treated with Mabisi, FOS, and sterile water.

## 4.0 Discussion

This study aimed to evaluate the capacity of two foodborne pathogens to invade gut microbiota under various treatments. Of the two strains of pathogens added to colon digestion units, only *Escherichia coli* established in the gut microbiota treated with FOS, however not in Mabisi and sterile water treatments.

*Escherichia coli* only established in the gut microbiota when exposed to FOS of all treatments, suggesting Mabisi treatment may prevent proliferation of added pathogens in the gut microbiota. Some pathogenic *Escherichia coli* strains are reported to survive *in vitro* human digestion stress. In a study by Buberg and colleagues^17^, 31 *E. coli* isolates were digested and all survived with 24 isolates able to transfer antibiotic resistant gene (*bla_CMY2_)* located on its plasmids. In our case, the non-virulent *E. coli* strain (K12) exposed to Mabisi treated gut microbiota could have suffered competition from commensal bacteria, for instance *Pediococcus,* explaining its failure to establish.

*Pediococcus* and others in the gut microbiota are lactic and acetic acid producing with capabilities of decreasing stool pH therefore creating unsuitable environment for pathogens to thrive. In line with our finding, a study investigating the probiotic effects of *Pediococcus pentosaceus L1* on enterotoxigenic *Escherichia coli* (ETEC) F4^+^ found that *P. pentosaceus L1* thrived at a low pH of 2.5, thereby inhibiting the growth of ETEC F4^+31^. A study to establish threshold concentration of Mabisi that could prevent virulent

*Escherichia coli* is relevant for validation and establishing efficacy to prevent virulent *Escherichia coli*. Leale and colleagues did not find an invader (e coli) following repeated challenges (100 generations) of natural microbial communiteis^32^. They attribute the failure of invasion of natural microbial community to low pH as main driving factor preventing invasion by *E. coli*.

Despite adding a high concentration of live non-virulent model pathogenic strains, *Listeria innocua* at about 5log_10_ cfu/ml summing up to a third of total by volume of the digestion content, we did not detect *Listeria innocua* in Mabisi, FOS, and Sterile water treated gut microbiota. The failure to trace the *Listeria innocua* could have been possibly attributed to multiple factors. Firstly, we used a simple digestion model, that did not include the mucosa layer to allow for *Listeria innocua* to establish through infection of host cells therefore inducing its own uptake through internalin and accessing intracellularly using actin-based motility. In animals, by evading the cellular immune response, *Listeria innocua* is expected to spread from cell to cell^33^. Secondly, *Listeria innocua* may have failed to thrive in the stool culture due to insufficient nutrients to support its growth as supported by the ecological principle of competitive exclusion. The commensal bacteria in the pooled stool culture could be competitively fitter compared to the non-virulent *Listeria innocua* strain used in our experiment. *Listeria innocua* and *Listeria monocytogenes* as demonstrated by McLaughlin et al in an *in-vitro* study declined survival rates in soil samples possibly due to presence of soil microbiota^34^.

Thirdly, nutrient starvation combined with decreased O_2_ concentration in an anaerobic setting could have also caused the failure for *Listeria innocua* to grow and invade. We are not surprised not finding the invader, Listeria because of low pH associated Mabisi that could prevent its growth. For the future, we recommend digestion models that account for mucosa layer to be suitable for studying *Listeria* invasion of the gut microbiota.

The clear separation observed among the Mabisi, FOS, and Sterile water treated gut microbiota in the PCoA plot (Figure 3A) suggests their microbial composition is distinguishable and that FOS addition has the strongest effect since FOS from the other treatments separates along the first axis and Mabisi and sterile water separate along the second axis. Addition of pathogen separates from the pathogen free treatments along the third axis, highlighting the detectable effect of pathogen exposure (see Figure 3B). In total, these 3 axes capture over 75% of variation. Mabisi efficacy on gut microbiota is important information as it may stimulate clinical context use to prevent and possibly treat specific gut microbiota imbalances. Similar TFF, particularly of dairy origin have demonstrated their potential to positively impact gut microbiota. A dairy based TFF reported to influence gut microbiota is Kefir with evidence to modulate gut microbiota to a structure that is critical in preventing obesity, type 2 diabetes mellitus, liver diseases, cardiovascular disorders and improve immune response^36^. Important components of TFF include peptides, bioactive compounds, and specific lactic acid producing (LAB) bacterial strains. Discriminant bacterial strain in Mabisi treatment is *Pediococcus*, when compared to Sterile water. These bacterial genera are known producers of metabolites such as SCFA that suppress proliferation of resident and foreign invading pathogens^37,38^. Results of this study consolidates the effect of Mabisi on gut microbiota reflecting on our previous evidence^39^ in which *Pediococcus* increased in Mabisi treated gut microbiota.

Our study further demonstrates that Mabisi is a major driver of SCFA production. Here, acetate, propionate, lactate, isobutyrate, formate and succinate were significantly in higher concentration in Mabisi treatment compared to positive control (FOS) and negative control. Experiment design using 5% FOS is a standard or moderate level of prebiotics that provide benefits without causing any risk to the existence of gut microbiota^40^. Despites use of 5% FOS, Mabisi produced more SCFA compared to FOS. Butyric acid a target SCFA monitored in our experiment was not detected. Low relative abundances of *Clostridium*, *Ruminococcus* and *Roseburia* species that are important producers of Butyric acid could have caused this very low production of butyric acid to undetectable levels^5^. Another factor would be to extend incubation period as production of butyric acid may occur later during fermentation^10^. Confirmation of Mabisi causing increase in production of SCFA is important considering its consumption by children. Higher production of SCFA on exposure to Mabisi indicates its potential as a prebiotic or probiotic agent. An influx of SCFA promotes health of gut epithelia cells and growth of commensal bacteria^41–46^.

Studying properties of TFF or other food material using in vitro approaches is ethically recommended prior to clinical trials. The INFOGEST static digestion and digestion models in general are used successfully to study food properties. This model is reproducible and allows for a wider range of technical replicates. Another fact is the consensus among Food digestion scientists in accepting its outcomes making this study valid and comparable to similar studies. Lastly, the model is cheap and easy to implement, however, it may not depict the exact human physiological processes when compared to more complex digestion models such as the MiGUT^47^. Current evidence of beneficial effect of Mabisi on gut microbiota is now based on the static INFOGEST digestion and fermentation and the SHIME model. The work Alekseeva and colleagues conducted (in press) using the SHIME model on effect of Mabisi show similar properties of increasing SCFA besides modulating *in vitro* gut microbiota with an abundance of beneficial bacteria^48^. With the current evidence on Mabisi, our next steps are to consolidate evidence on prebiotic and probiotic effect of Mabisi and related TFF that may create the need to conduct an intervention study. Firstly, Mabisi could be studied further for its prebiotic effect on a wide range of gut microbiota using more complex dynamic models that allow for more replications compared to the SHIME model. Another area of interest could be isolation of *Pediococcus*, bacteria that stand out on exposure to Mabisi and investigate it further for its functional genes and physicochemical properties.

Mabisi used here prevented *E.coli* K12 invasion by modulating the gut microbiota, however, more evidence is needed for its efficacy against enterotoxigenic *E coli* strains and other similar pathogens. For the future, a dynamic *in vitro* model could be used to study pathogen invasion and modulation of gut microbiota using multiple TFF sources with known microbiota. This study has provided information on potential use of TFF as a preferred medium for production of therapeutic food for use in children especially in low-income environments. Outcomes of this study are of public health importance especially in the face of common foodborne outbreaks caused by Shiga toxin producing bacteria strain such as *Escherichia coli* 0157:H7^49^. Mabisi has shown potential for therapeutic support in managing *Escherichia coli* 0157:H7 infections, however, more evidence is needed that shows its efficacy to suppress a wide range of virulent *E coli* strains.

## Acknowledgements

We gratefully acknowledge the families in Zambia who provided stool samples for this work, and Dr. Alanna Leale and Dr Bwalya Katati for their help in implementing this experiment. Sibbe Bakker contributed to data analysis and visualization and led efforts to ensure the study data adhered to FAIR principles (Findable, Accessible, Interoperable, and Reusable). Finally, this work was made possible through funding from Nutricia foundation and the Wageningen Graduate Schools (WGS) PhD fellowship programme.

## 6.0 Author contribution

**MN:** Conceptualization, data curation, data analysis, visualization, project administration, writing (original manuscript), writing (editing and reviews).

**OM:** Data curation, data analysis, visualization, validation, writing (original manuscript).

**EM:** Conceptualization, project administration, writing (original manuscript), validation, writing (editing and reviews), supervision.

**JS:** project administration, writing (original manuscript), writing (editing and reviews), validation, supervision.

**BZ:** Conceptualization, data curation, data analysis, visualization, project administration, validation, writing (original manuscript), writing (editing and reviews), supervision.

**SS:** Conceptualization, data curation, data analysis, visualization, project administration, validation, writing (original manuscript), writing (editing and reviews), supervision.

## 9.0 Supplemental material

**Supplement Figure 1.**
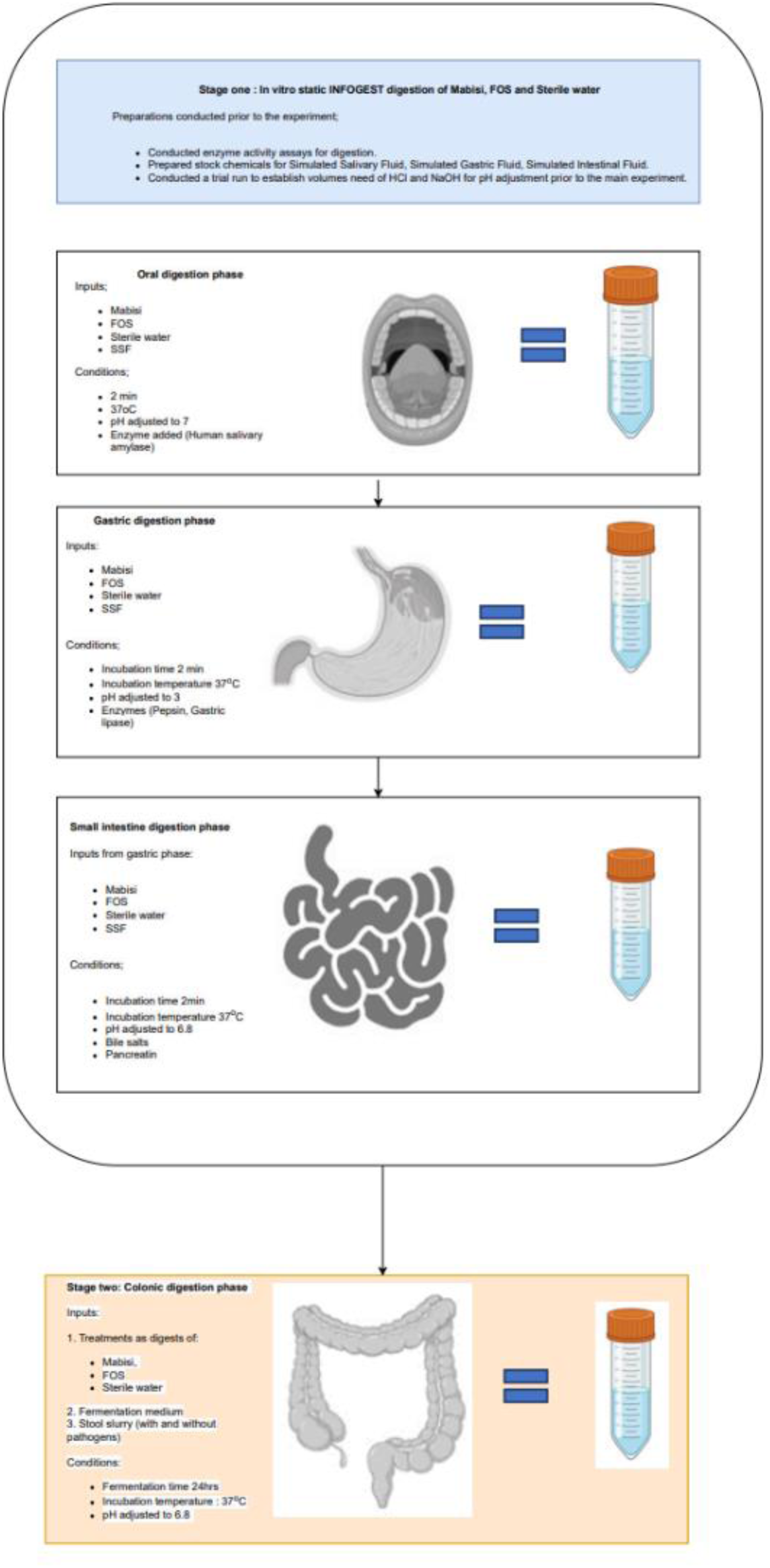
A flow chart showing in vitro static INFOGEST and colon digestion inputs, conditions, and treatments of experiment inputs and its conditions. In the second colon digestion, Mabisi, Fructooligosaccharides (FOS) and Sterile water digests are exposed to pooled stool slurry with or without pathogens.

**Supplement Figure 2.**
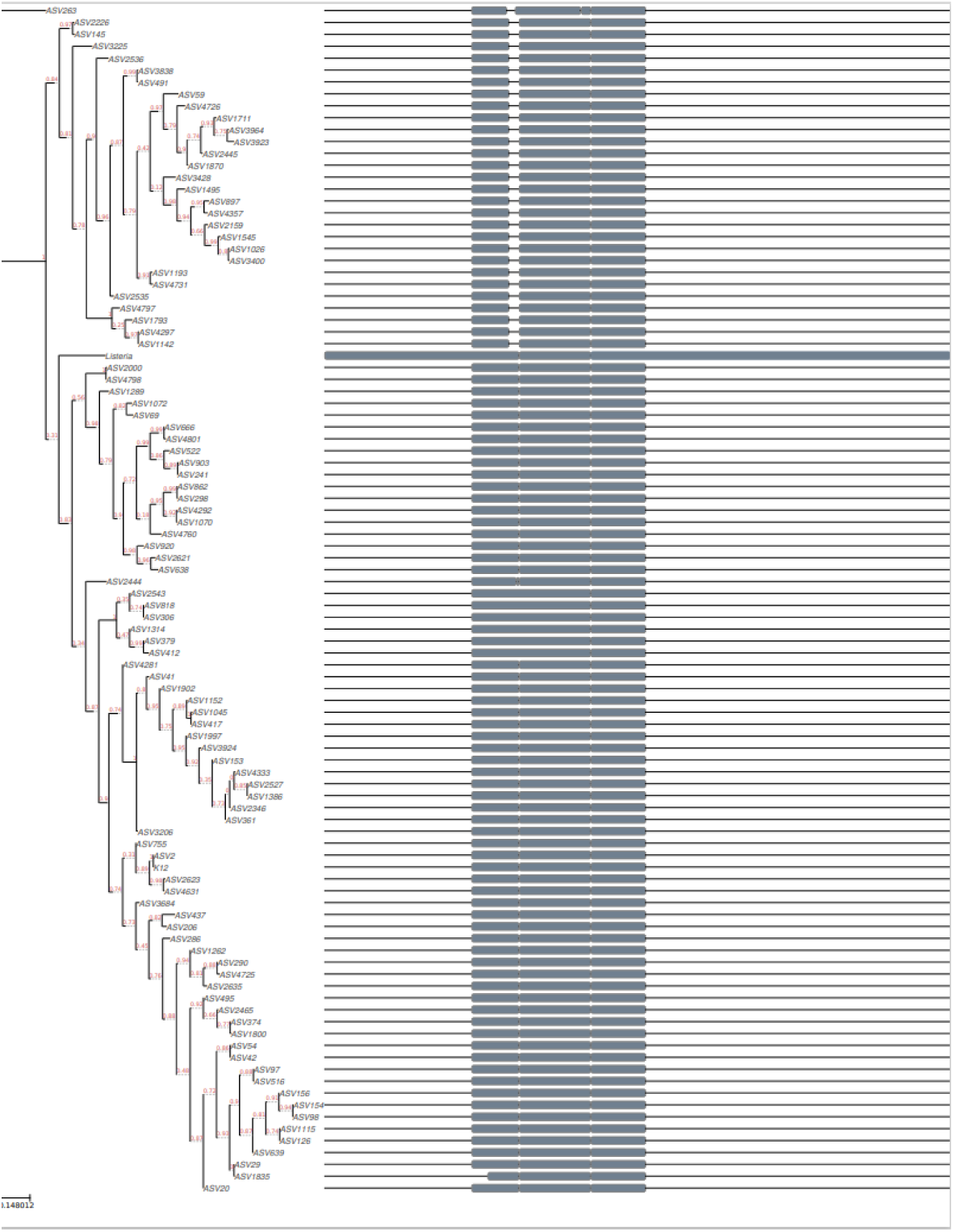
Shows phylogenetic tree of proteobacteria ASVs and gut microbiota pathogen added strains, *Escherichia coli K12* DSM 498 *and Listeria innocua ATCC 33090 DSM20649*.

**Supplement Figure 3.**
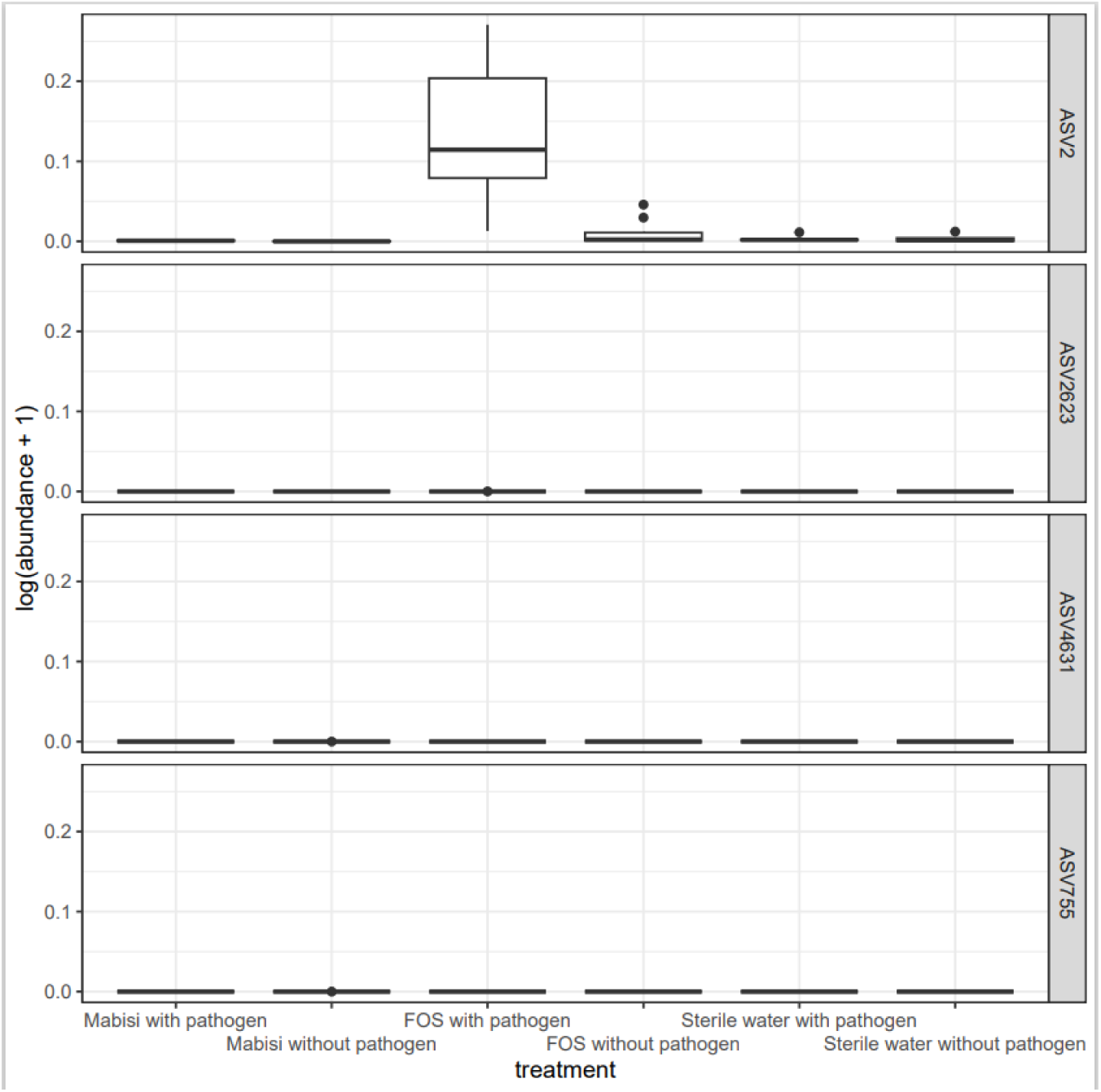
Boxplots showing log (absolute abundance + 1) transformed relative abundance of closely related ASVs to Escherichia coli K12 DSM 498.

**Supplement Figure 4.**
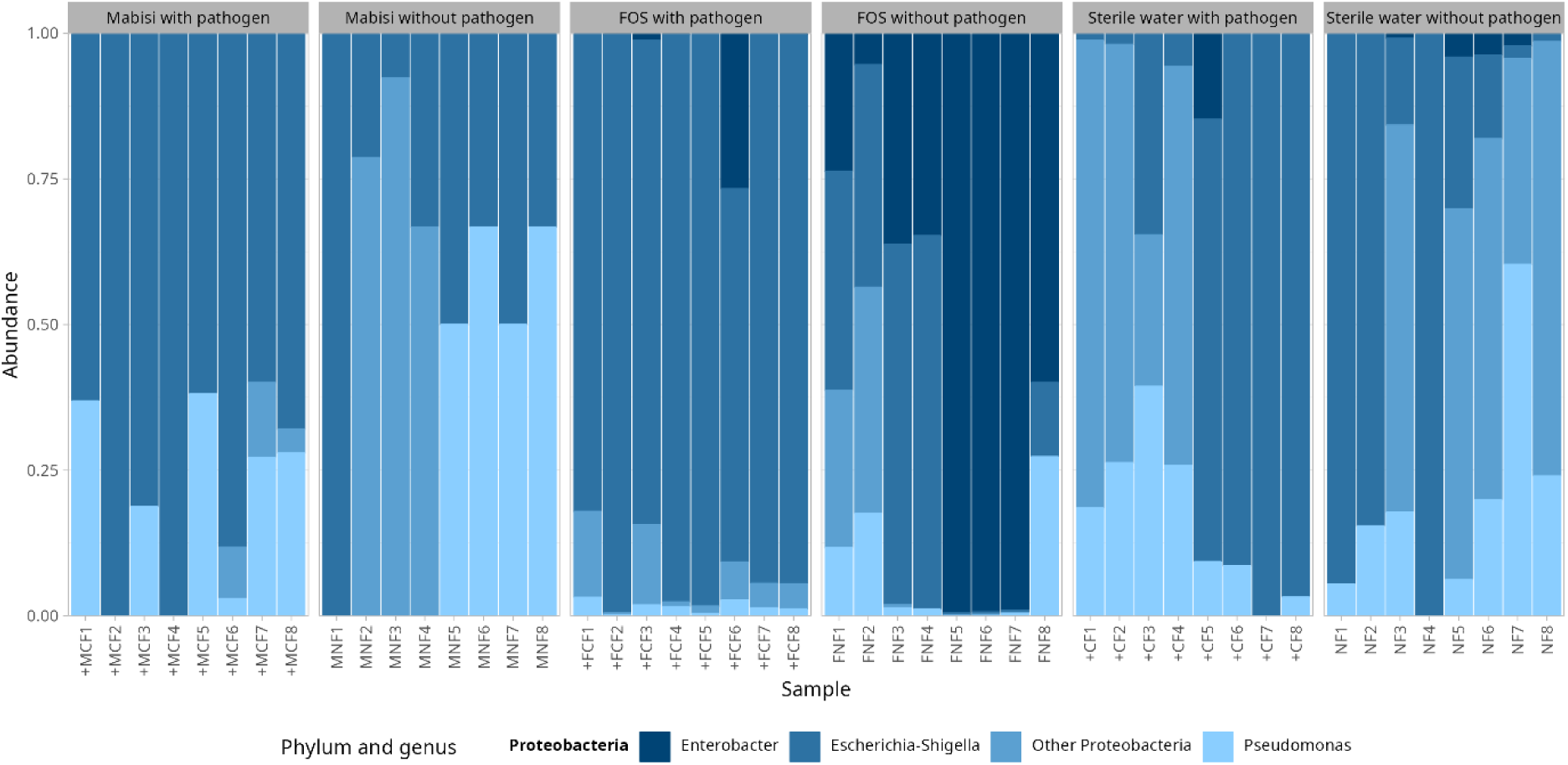
Relative abundance of top three proteobacteria after treatment with Mabisi, FOS, and sterile water. The plus (+) sign show fermentation units with pathogen added in each treatment.

**Supplement Figure 5.**
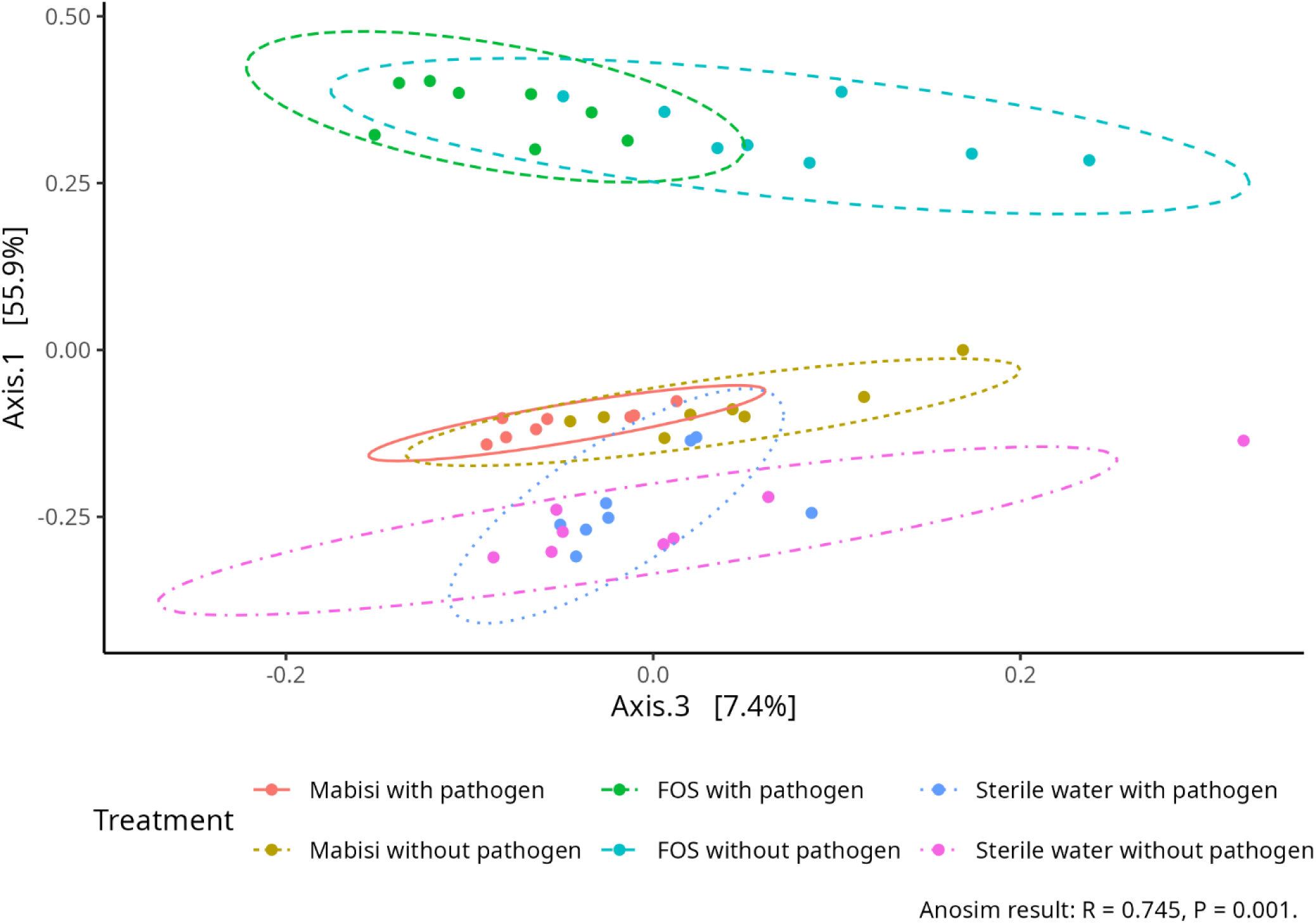
Principal Coordinate Analysis showing effect of addition of pathogens to treatments correlating to axis 3 with variation of 7.4% explained variation.

**Supplement Figure 6.**
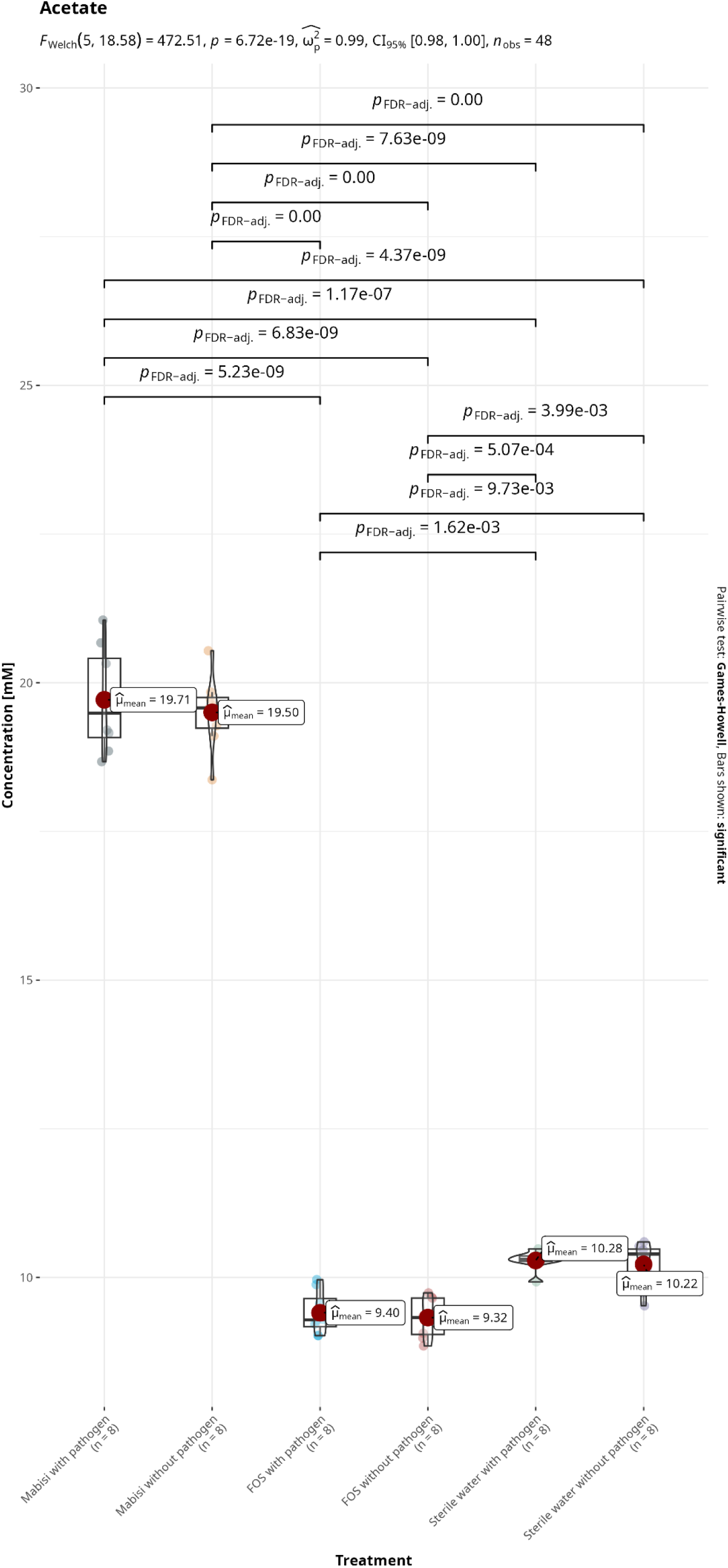
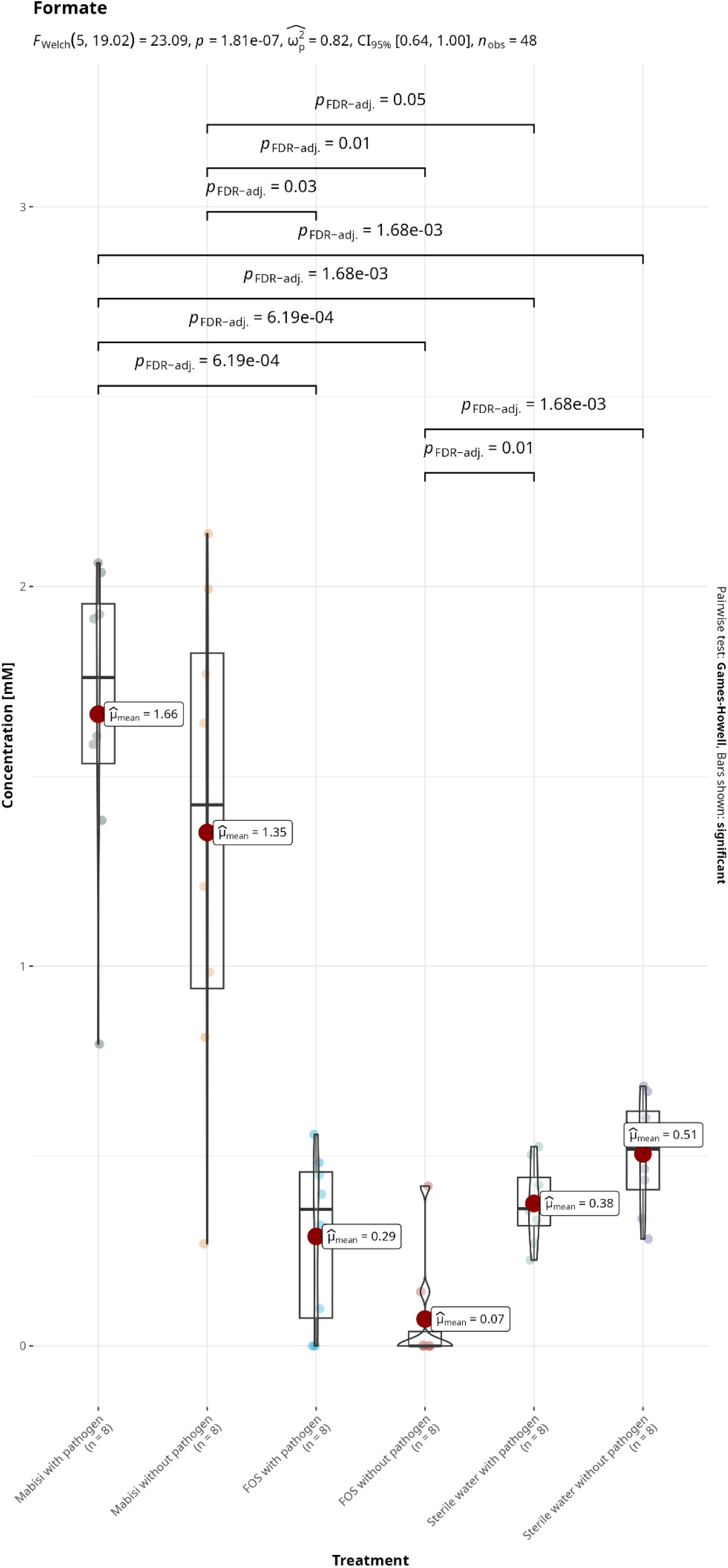
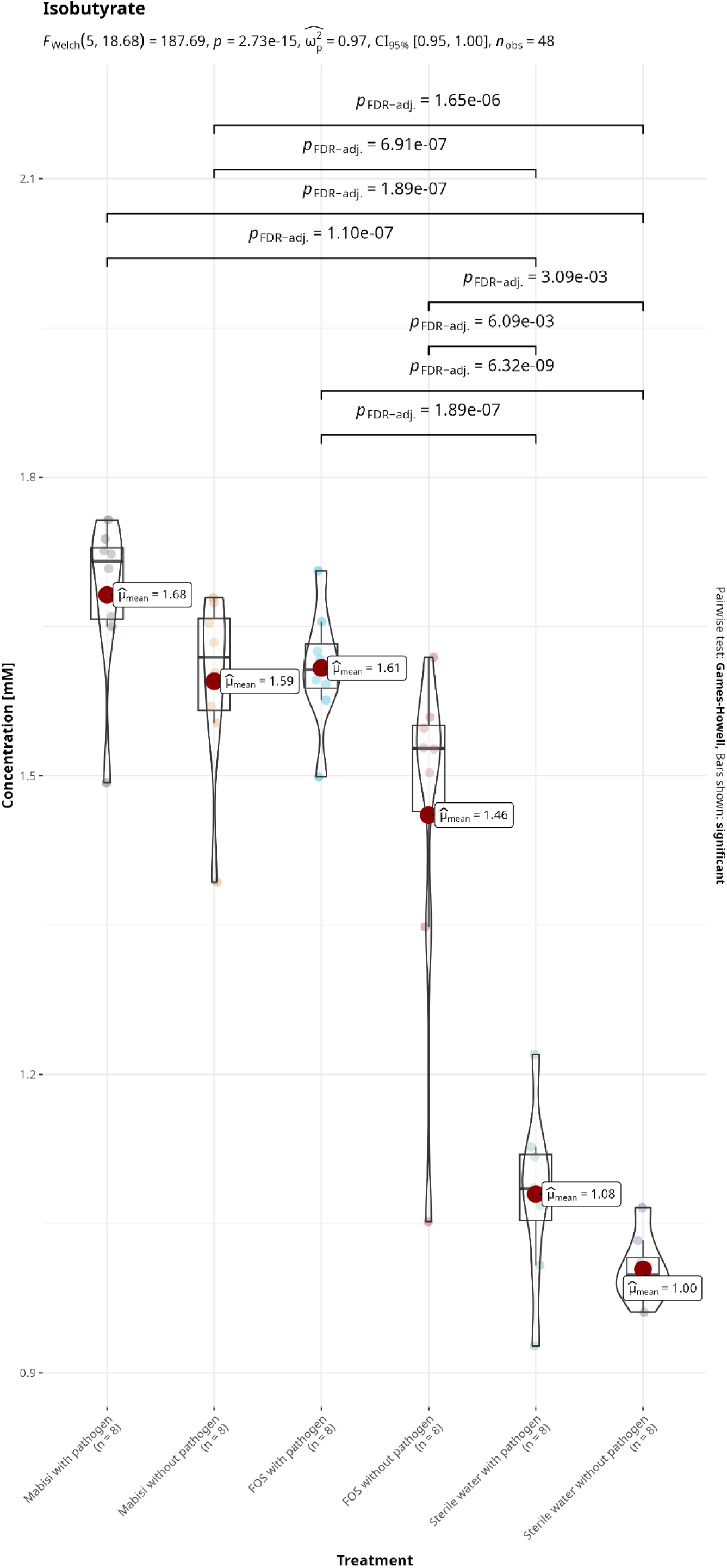
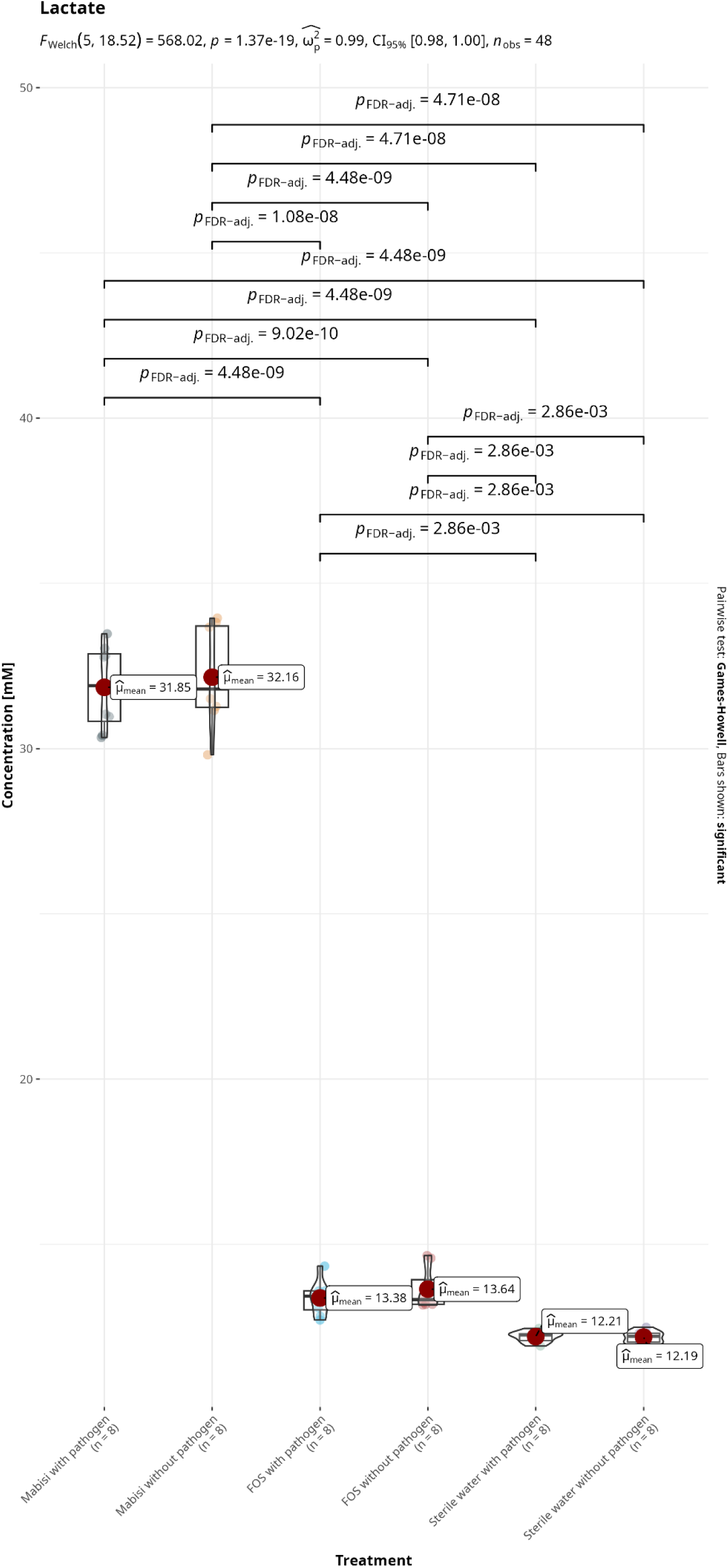
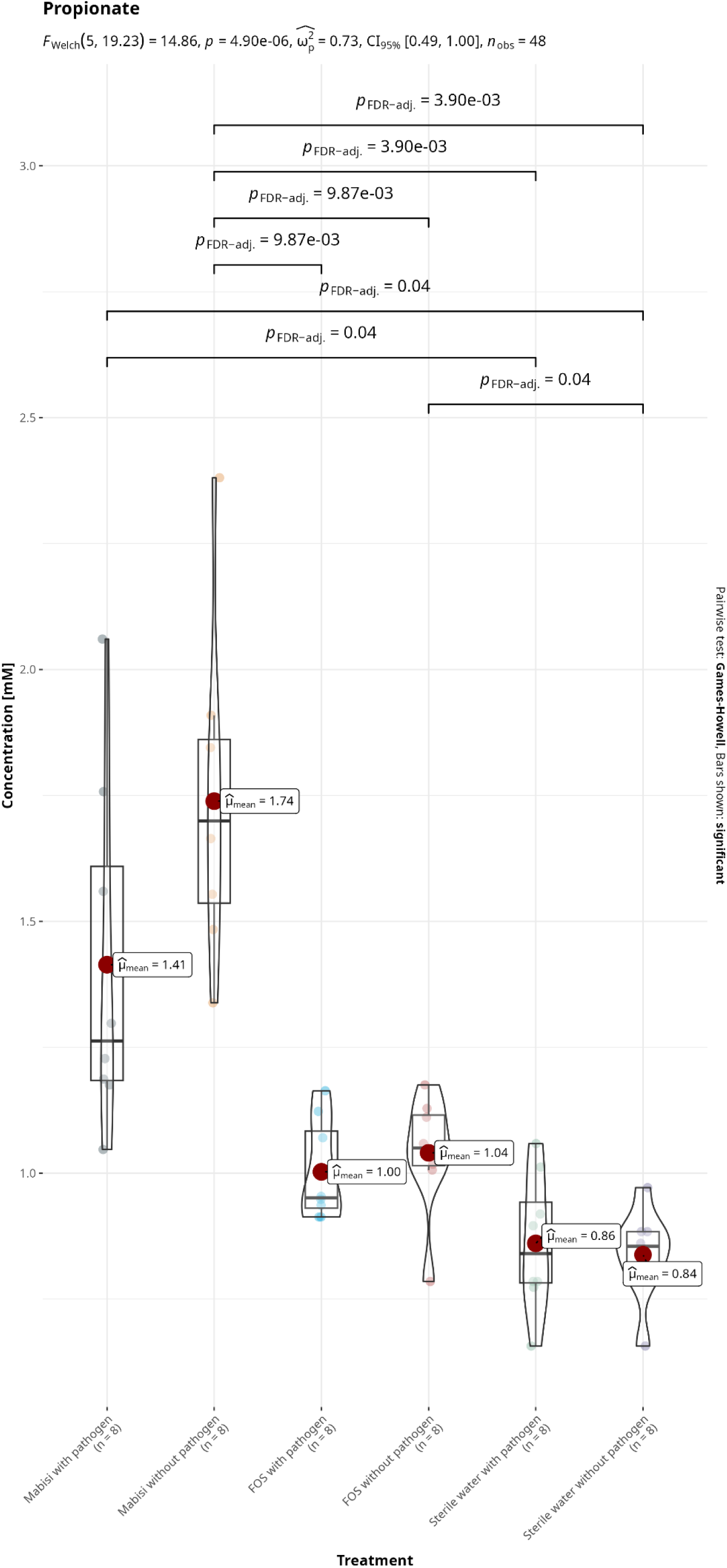
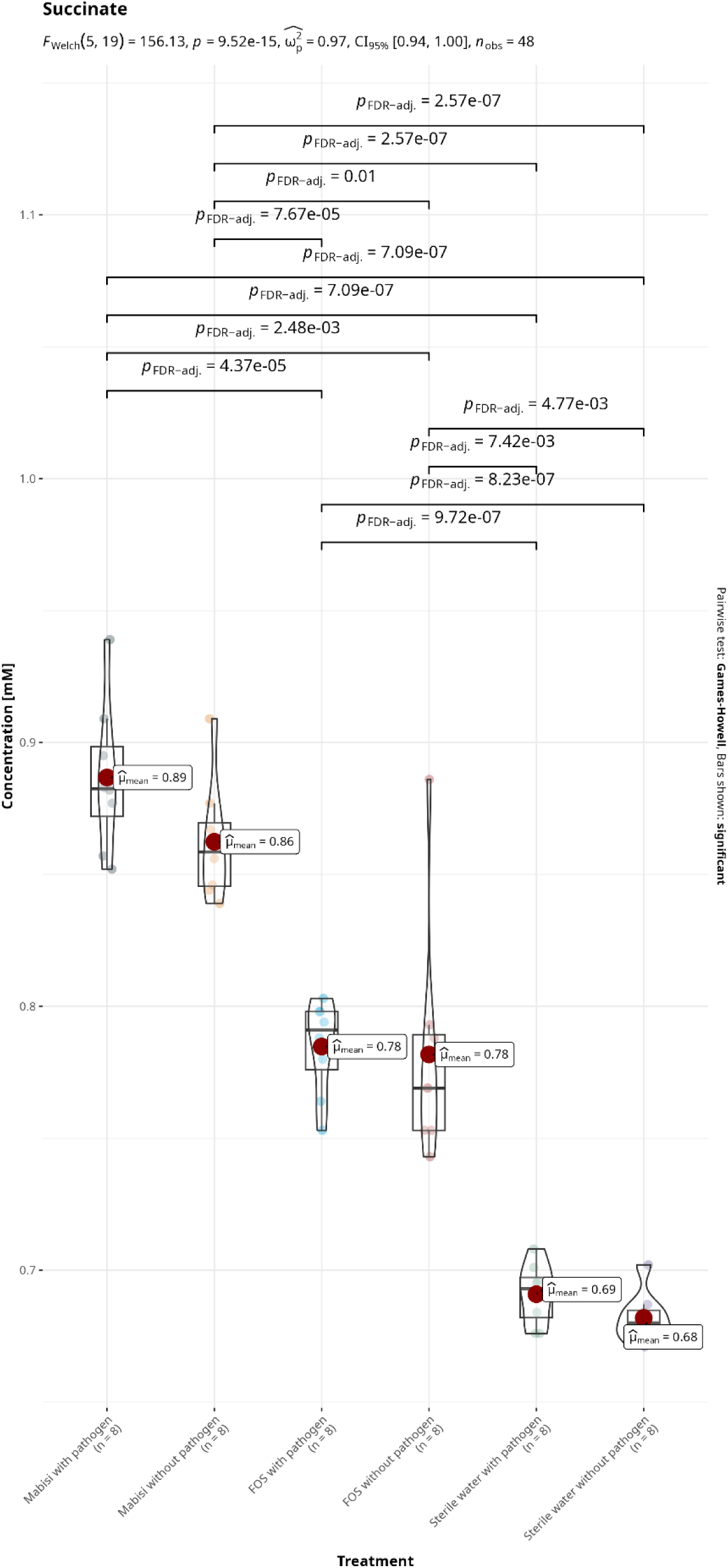
Short chain fatty acid (acetate, formate, lactate, propionate, isobutyrate and succinate) concentrations (mM) in colon digestion units treated with Mabisi, FOS, and sterile water. Treatments are compared for concentration using Games Howell pairwise test.

**Supplement Figure 7.**
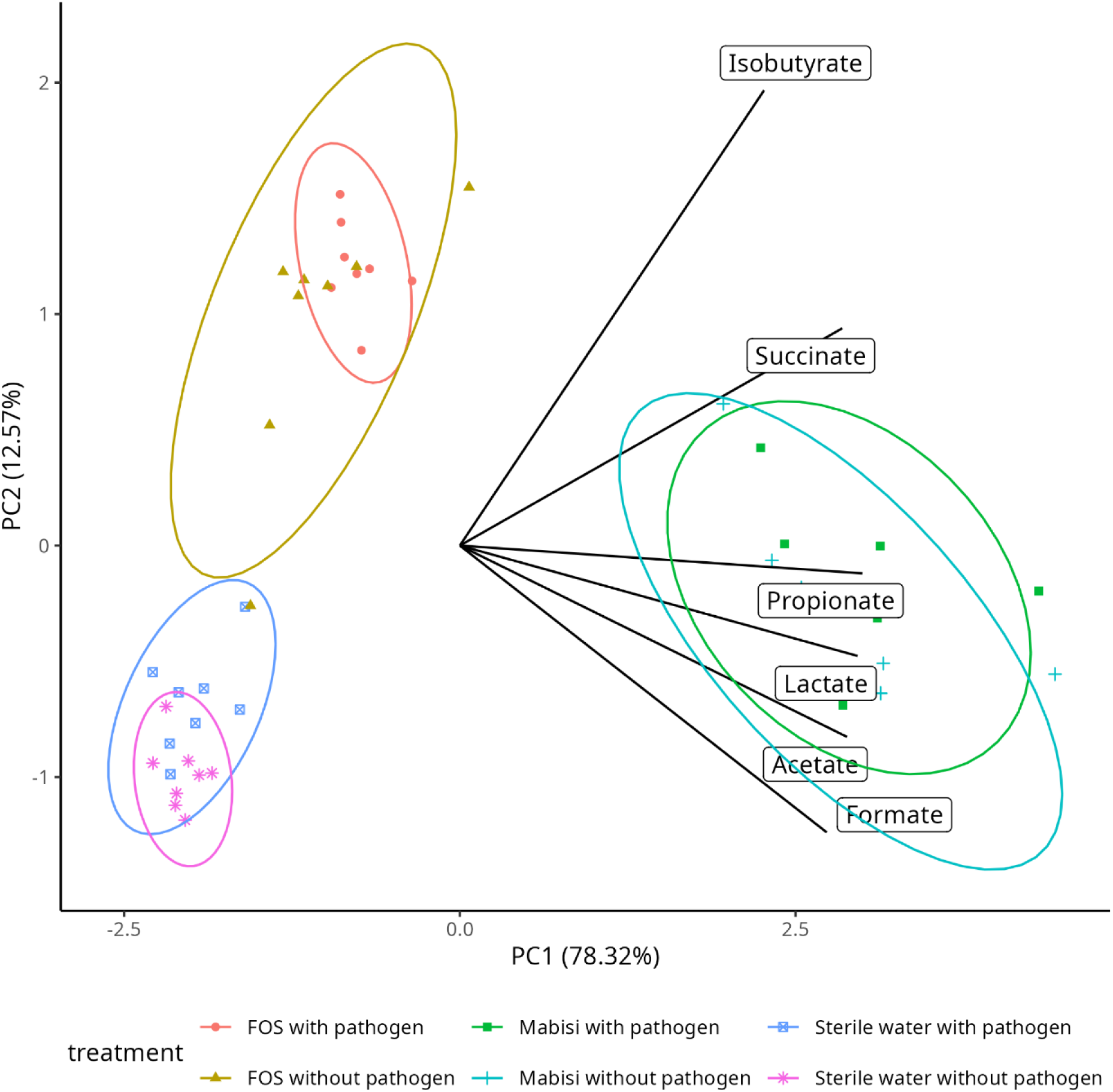
A PCA ordination of SCFA production of Mabisi, FOS and Sterile water treatments. The PCA reveal Mabisi treated colon digestion units share orthogonal direction for acetate, propionate, lactate, formate and succinate production compared to sterile water and FOS.

**Supplement Table 1.** Pairwise PERMANOVA analysis of treatments after exposure to Mabisi, FOS and sterile water (negative control).

| Pairs of treatments | Sum of squares | F.Model | R2 | p.value | p.adjusted |
| --- | --- | --- | --- | --- | --- |
| **Mabisi with pathogen vs Mabisi without pathogen** | **0.444** | **1.101** | **0.073** | **0.187** | **1.000** |
| Mabisi with pathogen vs Sterile water with pathogen | 0.729 | 1.703 | 0.108 | 0.000 | 0.003* |
| Mabisi with pathogen vs Sterile water without pathogen | 0.729 | 1.667 | 0.106 | 0.000 | 0.003* |
| Mabisi with pathogen vs FOS with pathogen | 0.874 | 2.104 | 0.131 | 0.000 | 0.003* |
| Mabisi with pathogen vs FOS without pathogen | 0.851 | 2.111 | 0.131 | 0.000 | 0.003* |
| Mabisi without pathogen vs Sterile water with pathogen | 0.773 | 1.856 | 0.117 | 0.000 | 0.003* |
| Mabisi without pathogen vs Sterile water without pathogen | 0.775 | 1.820 | 0.115 | 0.000 | 0.006* |
| Mabisi without pathogen vs FOS with pathogen | 0.985 | 2.437 | 0.148 | 0.000 | 0.003* |
| Mabisi without pathogen vs FOS without pathogen | 0.942 | 2.406 | 0.147 | 0.000 | 0.003* |
| **Sterile water with pathogen vs Sterile water without pathogen** | **0.511** | **1.136** | **0.075** | **0.055** | **0.831** |
| Sterile water with pathogen vs FOS with pathogen | 0.885 | 2.065 | 0.129 | 0.000 | 0.006* |
| Sterile water with pathogen vs FOS without pathogen | 0.924 | 2.223 | 0.137 | 0.000 | 0.003* |
| Sterile water without pathogen vs FOS with pathogen | 0.844 | 1.928 | 0.121 | 0.000 | 0.006* |
| Sterile water without pathogen vs FOS without pathogen | 0.888 | 2.089 | 0.130 | 0.001 | 0.009* |
| **FOS with pathogen vs FOS without pathogen** | **0.668** | **1.657** | **0.106** | **0.003** | **0.048\*** |

Asterix sign (\*) indicate statistical significance at $p < 0.05$ .

**Supplement Table 2.** Results of PERMANOVA analysis of fermentation units (gut microbiota).

| **Pairs of fermentation units** | **Df** | **Sum of Squares** | **F.Model** | **R2** | **p.value** | **p.adjusted** |
| --- | --- | --- | --- | --- | --- | --- |
| Mabisi vs Negative control | 1 | 1.03 | 2.38 | 0.07 | 0.000 | 0.001* |
| Mabisi vs FOS | 1 | 1.27 | 3.07 | 0.09 | 0.000 | 0.001* |
| Negative control vs FOS | 1 | 1.18 | 2.70 | 0.08 | 0.000 | 0.001* |

Asterix sign (\*) indicate statistical significance at $p < 0.05$ .

